# Comparative Genomics and Directed Evolution Reveal Genetic Determinants of Extreme UVC Radiation Tolerance in Bacteria Recovered from the Stratosphere

**DOI:** 10.1101/2023.03.27.534493

**Authors:** Adam J. Ellington, Tyler J. Schult, Christopher R. Reisch, Brent C. Christner

## Abstract

Aerosolized microbes surviving transport to and in the stratosphere endure extremes of low temperature, atmospheric pressure, and relative humidity, and high shortwave ultraviolet radiation flux. However, the genetic determinants for traits enabling resistance to the combination of stresses experienced by microbes in the high atmosphere have not been systematically investigated. In this study, we examined Proteobacteria and Actinobacteria isolated from the stratosphere (18 to 29 km ASL) and that demonstrated high tolerance to desiccation (15-25% RH) and UVC radiation (UVCR; λ= 254 nm). Closely related reference strains were more sensitive to UVCR than the stratospheric isolates, indicating that extreme resistance is not universally distributed in these phylogenetically related bacteria. Comparative genomic analyses revealed DNA repair and antioxidant defense genes in the isolates that are not possessed by the related reference strains, including genes encoding photolyase, DNA nucleases and helicases, and catalases. Directed evolution by repeated exposure to increasing doses of UVCR improved the LD_90_ in a sensitive reference strain by ∼3.5-fold. The mutations acquired in *Curtobacterium flaccumfaciens pv. flaccumfaciens* strain DSM 20129 incrementally increased its UVCR resistance, with the accumulation of 20 point mutations in protein coding genes increasing tolerance to a level approaching that of stratospheric isolate *Curtobacterium* sp. L6-1. The genetic basis for the increased UVCR tolerance phenotypes observed is discussed, with a specific emphasis on the role of genes involved in DNA repair and detoxification of reactive oxygen species.

**Importance:** Ultraviolet radiation is omnipresent in sunlight and has important biological effects on organisms. The stratosphere is the only location on Earth where microbes receive natural exposure to highly mutagenic wavelengths (<280 nm) of ultraviolet radiation. Genetic studies of bacteria from an environment that selects for extreme ultraviolet radiation resistant phenotypes has expanded what is known from studies of model species (e.g., *E. coli*) and identified potentially novel protection and repair strategies. In addition to deepening understanding of ultraviolet radiation photobiology in atmospheric microbes and bacteria in general, these advancements are also highly relevant to astrobiology and space biology. The cold, dry, hypobaric, and high radiation environment of the stratosphere provides an earthly analog for thin extraterrestrial atmospheres (e.g., Mars) and is ideal for bioprospecting extremophile phenotypes that enable engineering of genetic stability and functionality in bio-based space life-support systems or any application where long-term persistence is desirable (e.g., biocontrol).

## Introduction

The persistence of microbes during atmospheric transport and deposition has important implications to human health (1, 2), agriculture (3–5), meteorology (6–8), and astrobiology (9, 10). Many studies have focused on microbial aerosols in the lowest layer of the troposphere, where the air is in contact with and affected by the surface (i.e., convective boundary layer, CBL). However, the composition of microbial assemblages in the CBL can be quite different from that of overlying air masses in the ‘free troposphere’ (11, 12). Importantly, environmental conditions in the CBL are unrepresentative of those at higher altitudes, where biological stresses (e.g., water loss and ultraviolet radiation, UVR) become increasingly intense. Given that 10^23^ to 10^24^ microbes are estimated to disperse annually in the Earth-atmosphere system (13), new information is needed to determine the genetic basis and mechanisms enabling survival under the extreme conditions associated with atmospheric transport.

The stratosphere is the atmospheric layer above the troposphere (∼15 to 50 km above sea level, ASL) and is characterized by low temperature, atmospheric pressure, and relative humidity (RH) and high UVR flux (9, 12–16). This combination of conditions is unique to any other location on Earth and similar to conditions on the surface of Mars (14, 15), making the stratosphere a relevant astrobiological analog to examine microbial survival potential on alien worlds (16–19). Microbes that survive conditions in the stratosphere are limited to those tolerant of water loss and exposure to high levels of UVR (20). The UVR spectrum consists of three types: UVAR (320-400 nm), UVBR (280-320 nm), and UVCR (100-280 nm). Oxygen and ozone in the lower stratosphere strongly absorb short wavelength UVR, and consequently, UVCR is not observed at lower altitudes in the troposphere (21). As such, the stratosphere is the only natural laboratory on Earth to examine the biological effects of high energy UVR wavelengths.

UVR damages cells via direct and indirect effects. Direct absorption of UVR by DNA forms pyrimidine dimers that inhibit transcription and replication and can lead to single-stranded breaks in the sugar-phosphate backbone (22). Indirect effects of UVR occur through the generation of reactive oxygen species (ROS), which can induce single- and double-strand breaks, apurinic sites, and base damage in DNA, as well as the oxidation and chemical modification of proteins and lipids (23). Bacterial tolerance to UV is conferred by a complex network of genetic and physiological systems termed the “UV-resistome”, which has five basic functions: sense, shield, detoxify, repair, and tolerate (24). Photoreceptors and other stress sensors <u>sense</u> specific UV wavelengths and DNA damage, triggering a regulatory cascade that elicits specific cellular responses (25, 26). Specialized membrane proteins and UV-absorbing pigments <u>shield</u> intracellular components from irradiation (27). ROS-scavengers <u>detoxify</u> reactive species to prevent oxidative damage and maintain cellular redox homeostasis (28). Efficient DNA repair proteins <u>repair</u> genetic damages (29). And finally, error-prone polymerases <u>tolerate</u> and bypass unrepaired lesions to ensure survival of the cell at the cost of introducing mutations to the genome (30). The general response of bacteria to UVR and the DNA repair strategies utilized have been well-studied in organisms such as *E. coli* (22, 29, 31). However, model species are not representative of natural bacterial populations, and in particular, those that disseminate widely in the atmosphere.

Previously, we sampled aerosols to altitudes of 38 km ASL using a specialized payload attached to a helium balloon (32), isolated a variety of bacteria from the samples, and demonstrated their high tolerance to desiccation (15-25% RH) and UVCR (λ= 254 nm) (20).

Several of the isolates displayed levels of resistance rivaling that of the highly radio-and xero-tolerant bacterium *Deinococcus radiodurans*. In this study, we sequenced the genomes of two highly tolerant isolates (*Curtobacterium* sp. L6-1 and *Noviherbaspirillum* sp. L7-7A) and used comparative genomic analyses with closely related, UV-sensitive reference strains to identify genetic features likely contributing to UVCR tolerance. We also used directed evolution of the UV-sensitive *Curtobacterium flaccumfaciens pv. flaccumfaciens* strain DSM 20129 (referred to hereafter as DSM 20129) to improve its UVCR tolerance and identified the mutations responsible for enhanced resistance. Additionally, we examined cellular ROS levels in the strains after UVCR exposure and assessed photolyase activities via an *in vivo* photorepair assay. Our results improve understanding of the mechanisms enabling bacterial survival during atmospheric transport and genetic determinants of UVR tolerance in taxa for which extreme resistance has not been previously reported or investigated.

## Results

### UVCR-Resistance in Stratospheric Isolates and Related Strains

Based on comparison of 16S rRNA gene sequences from strains L6-1 and L7-7A, these isolates were identified as members of the Gram-positive actinobacterial genus *Curtobacterium* and the Gram-negative betaproteobacterial genus *Noviherbaspirillum*, respectively (20). To assess whether the high UVCR tolerance observed in the isolates is a feature shared with other closely related taxa, actinobacterial and betaproteobacterial reference strains were identified (Fig. 1A), obtained from culture collections (Table A1), and screened for UVCR tolerance. The inactivation rate (*k*) and LD_90_ were derived from a survival curve for each strain (Table A2). *Curtobacterium* sp. L6-1 has higher UVCR tolerance (LD_90_ of 470 J m^-2^) than DSM 20129 (LD_90_ of 98 J m^-2^; Fig. 1B; Table S2). Similarly, *Noviherbasprillum* sp. L7-7A also displayed higher UVCR tolerance than the reference strains tested (Fig. 1C), with an average LD_90_ of 290 J m^-2^ compared to 195 J m^-2^ for *N. soli* SUEMI10, 63 J m^-2^ for *N. autotrophicum* TSA66, and 35 J m^-2^ for *N. denitrificans* TSA40 (Table A2).

**Fig. 1.**
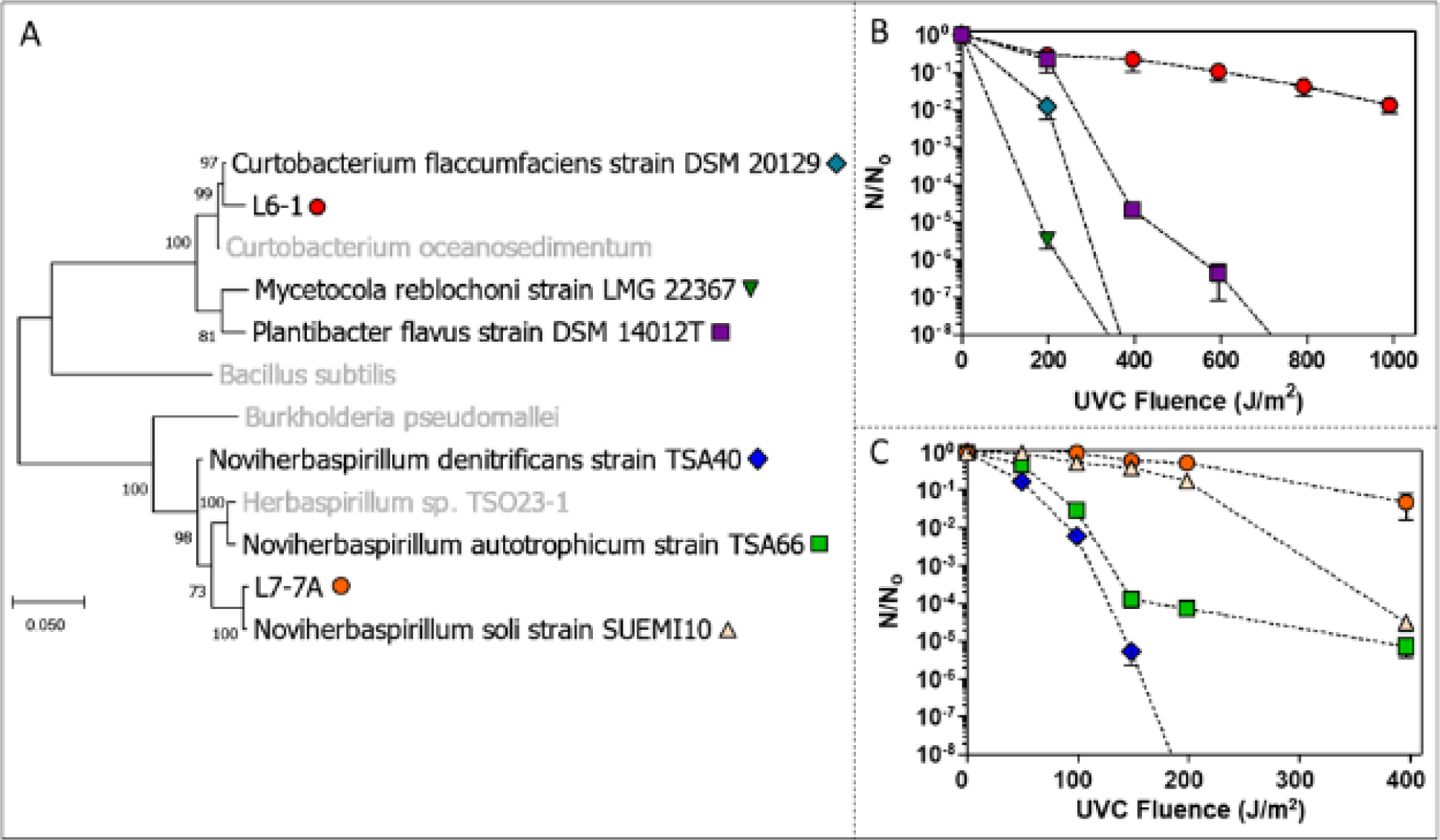
Comparison of phylogenetic relatedness among the bacterial strains and their tolerance to UVCR. **(A)** Maximum-likelihood analysis of 16S rRNA gene sequences from the stratospheric isolates and closely related strains. UVCR tolerance of the **(B)** actinobacterial and **(C)** betaproteobacterial strains based on the surviving number of cells (N) at each UVCR dose divided by that in unexposed populations (N_0_). The values plotted are averages from three independent replicates. Error bars represent SEM.

### Genetic Differences Between UVCR-Tolerant and -Sensitive Strains

The disparities in UVCR tolerance between the stratospheric isolates and reference strains suggested that L6-1 and L7-7A possess genetically discernable features contributing to this phenotype. To enable comparative genomic analysis, whole genome sequences were obtained for the *Curtobacterium* and *Noviherbaspirillum* strains and deposited in GenBank (33).

Strain L6-1 has a single 3.4 Mbp circular chromosome with a GC content of 72.0% and 3,158 genes encoding 12 rRNAs, 46 tRNAs, and 3,072 proteins (Table 1). Strain DSM 20129 possesses a circular chromosome and plasmid that together total 3.8 Mbp, with a GC content of 70.9%, and 3,616 genes encoding 9 rRNAs, 47 tRNAs, and 3,517 proteins. The two strains share 83% average nucleotide identity (ANI), which is below >95% values typically observed for closely related populations in a species taxon (34–37). For the *Noviherbaspirillum* strains, L7-7A possesses three circular chromosomes consisting of 5.2 Mbp and a GC content of 62.0%, with 4,771 genes encoding 15 rRNAs, 64 tRNAs, and 4,581 proteins. Although L7-7A has a genome size and content more similar to that of strain TSA66, it shares 79% ANI with both TSA66 and TSA40 and is likely a separate species (Table 1).

**Table 1:**
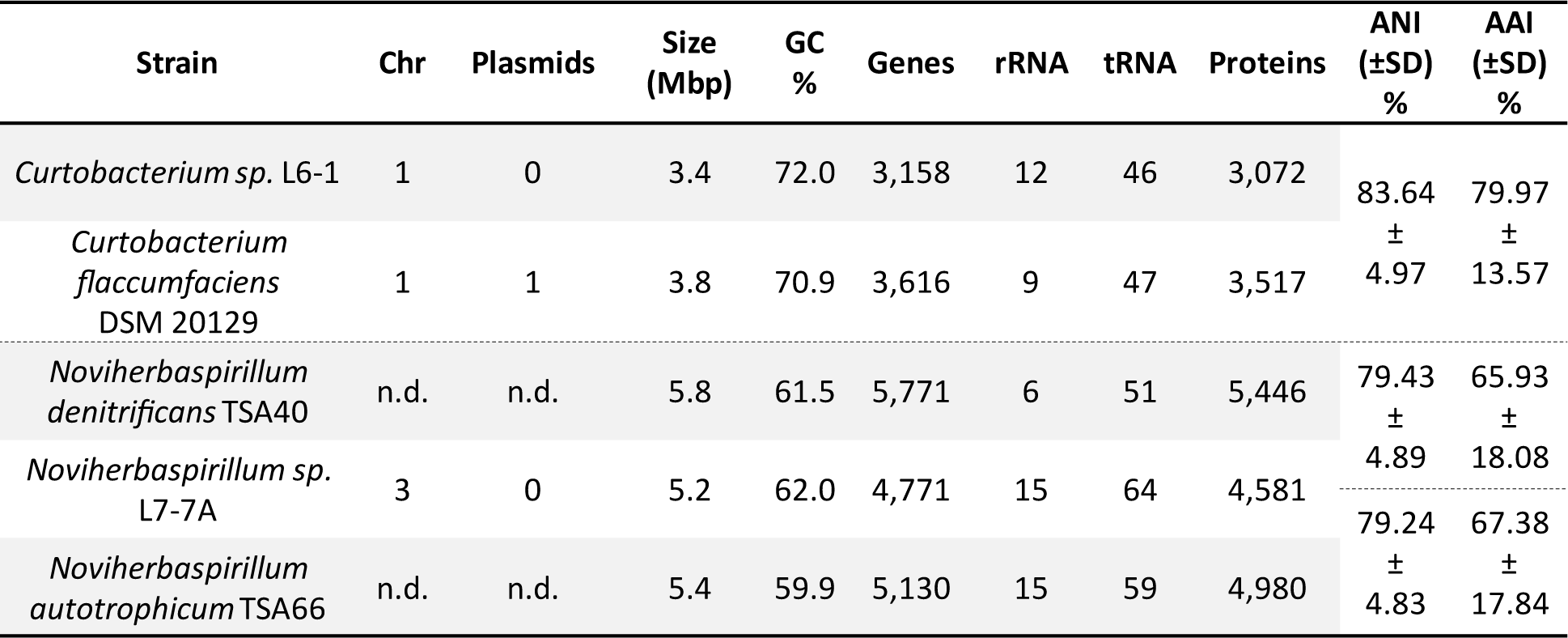
General genome features of the stratospheric isolates and closely related reference strains.

To analyze DNA repair and ROS detoxification pathways of the UV-resistome, genome comparisons using the RAST annotation server and the BLAST alignment tool were performed (Tables A3 and A4 for L6-1; Tables A5 and A6 for L7-7A; respectively). L6-1 encodes four additional proteins associated with DNA repair that are not found in DSM 20129 (Fig. 2; Table A3), including endonuclease VIII (*nei2*), two homologs of the UvrD/PcrA DNA helicases (*uvrD2* & *pcrA*), and a cryptochrome/photolyase family protein (*phr1*). In addition to DNA repair genes, L6-1 encodes a homolog of the KatA catalase (*katA*; Fig. 2; Table A4), the major catalase expressed in vegetative *Bacillus* (38), whereas DSM 20129 encodes a KatX homolog (Table A4), which has been demonstrated to protect germinating endospores from H_2_O_2_ stress (39). A homolog of the peroxide operon regulator PerR (*perR*) from *B. subtilis* is also found in L6-1, but not DSM 20129 (Fig. 2; Table A4). L6-1 also possesses an extra copy of the glutaredoxin-like protein NrdH (*nrdH2*; Fig. 2; Table A4) involved in disulfide reduction.

**Fig. 2.**
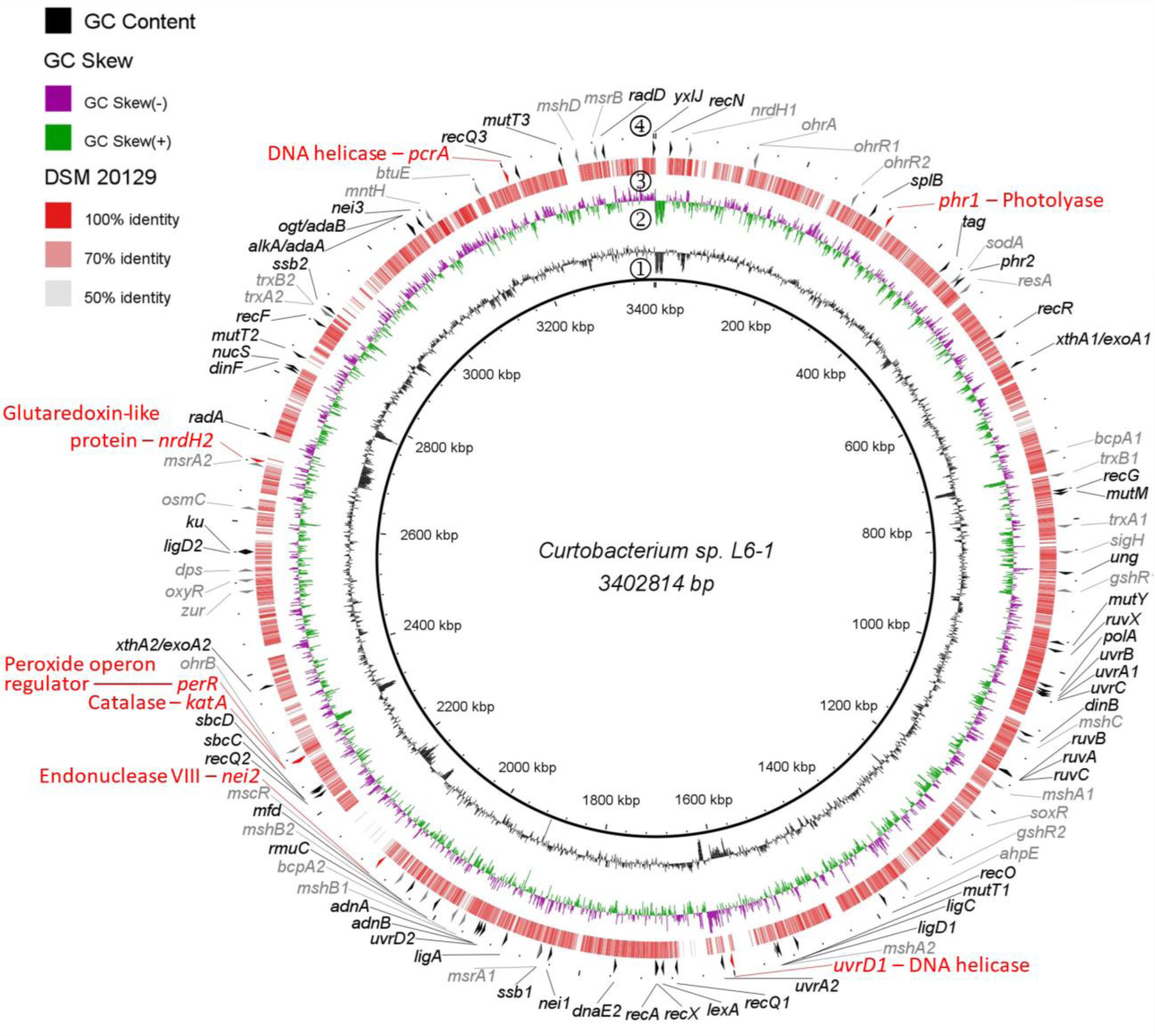
DNA repair and ROS detoxification genes in *Curtobacterium* sp. L6-1 not found in strain DSM 20129. Genomes were compared using the BLAST Ring Image Generator (BRIG) with the blastp algorithm. Rings represent the following (from innermost to outermost): **1)** GC Content, **2)** GC skew of the L6-1 genome, **3)** percent identity to L6-1 of protein orthologs found in strain DSM 20129, and **4)** DNA repair (**black**) and ROS detoxification (grey) genes. Genes present in L6-1 but absent in DSM 20129 are indicated in red text.

Of the nine DNA repair genes identified in L7-7A that are absent in strains TSA40 and TSA66 (Fig. 3; Table A5), there are additional copies of endonuclease III (*nth2* & *nth3*) and excinuclease UvrABC subunit A (*uvrA2*), along with the RecBCD nuclease subunit D (*recD*), DNA ligase C (*ligC*), alpha-ketoglutarate-dependent dioxygenase AlkB (*alkB*), error prone DNA polymerase V subunit D (*umuD*), and exonuclease SbcCD (*sbcC & sbcD*). Notably, all the *Noviherbaspirillum* strains lack the *recB* and *recC* genes, as well as genes for the alternative repair protein RecF of the RecFOR recombination pathway (Table A5). In addition to LigC, other essential components of the bacterial non-homologous end joining (NHEJ) repair pathway (LigD and Ku) are present in strains L7-7A and TSA40 (Table A5). Antioxidant genes unique to L7-7A (Fig. 3; Table A6) include additional copies of catalase (*katE*), a divalent metal cation transporter (*mntH2*), and the paraquat-inducible protein A (*pqiA2*). All three *Noviherbaspirillum* strains encode an ortholog of the *B. subtilis* KatX catalase (Table A6). However, L7-7A also encodes an ortholog of the *E. coli* KatE catalase (Fig. 3; Table A6). Though strain TSA40 possesses two genes for MntH, *mntH2* has low similarity to the *mntH2* of L7-7A (Table A6) and may have a different function.

**Fig. 3.**
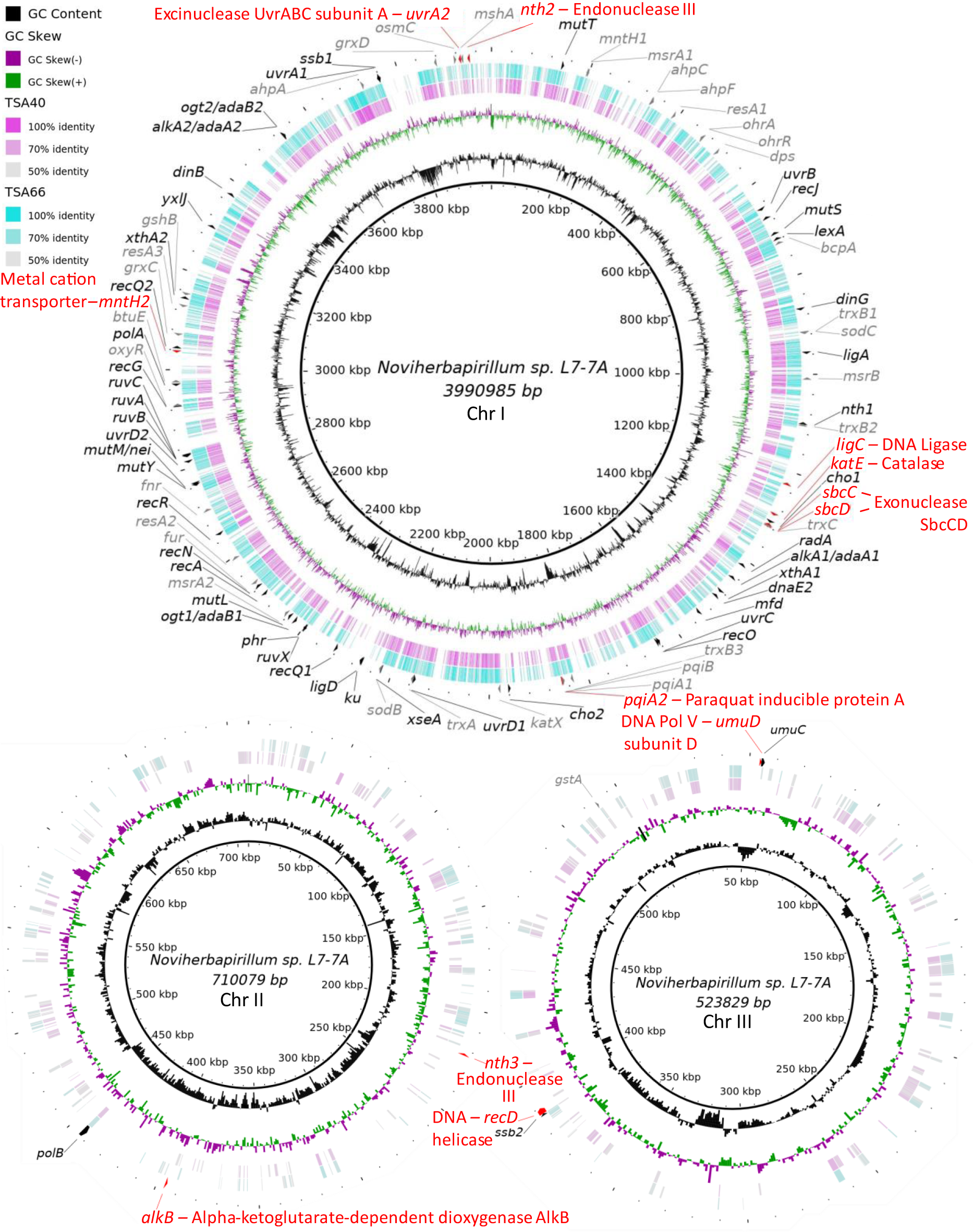
DNA repair and ROS detoxification genes in *Noviherbaspirillum* sp. L7-7A not found in strains TSA40 and TSA66. Genomes were compared using the BLAST Ring Image Generator (BRIG) with the blastp algorithm. Rings represent the following (from innermost to outermost): **1)** GC Content, **2)** GC skew of the L7-7A genome, **3)** percent identity to L7-7A of protein homologs found in strains TSA40 and TSA66, and **4)** DNA repair (**black**) and ROS detoxification (grey) genes. Genes present in L7-7A but absent in both TSA40 and TSA66 are indicated in red text.

### Directed Evolution of UVCR Tolerance in *C. flaccumfaciens* DSM 20129

Experiments designed to improve UVCR tolerance of the reference strains through repeated exposure to sublethal doses of UVCR (Fig. 4A) were partially successful. After 10 rounds of selection, we were unable to significantly improve the UVCR tolerance of the *Noviherbaspirillum* strains (data not shown). However, decreased UVCR sensitivity occurred incrementally in strain DSM 20129 during 12 rounds of selection, increasing its LD_90_ by 3.5-fold in comparison to the founder population and to a level comparable with isolate L6-1 (Fig. 4B).

**Fig. 4.**
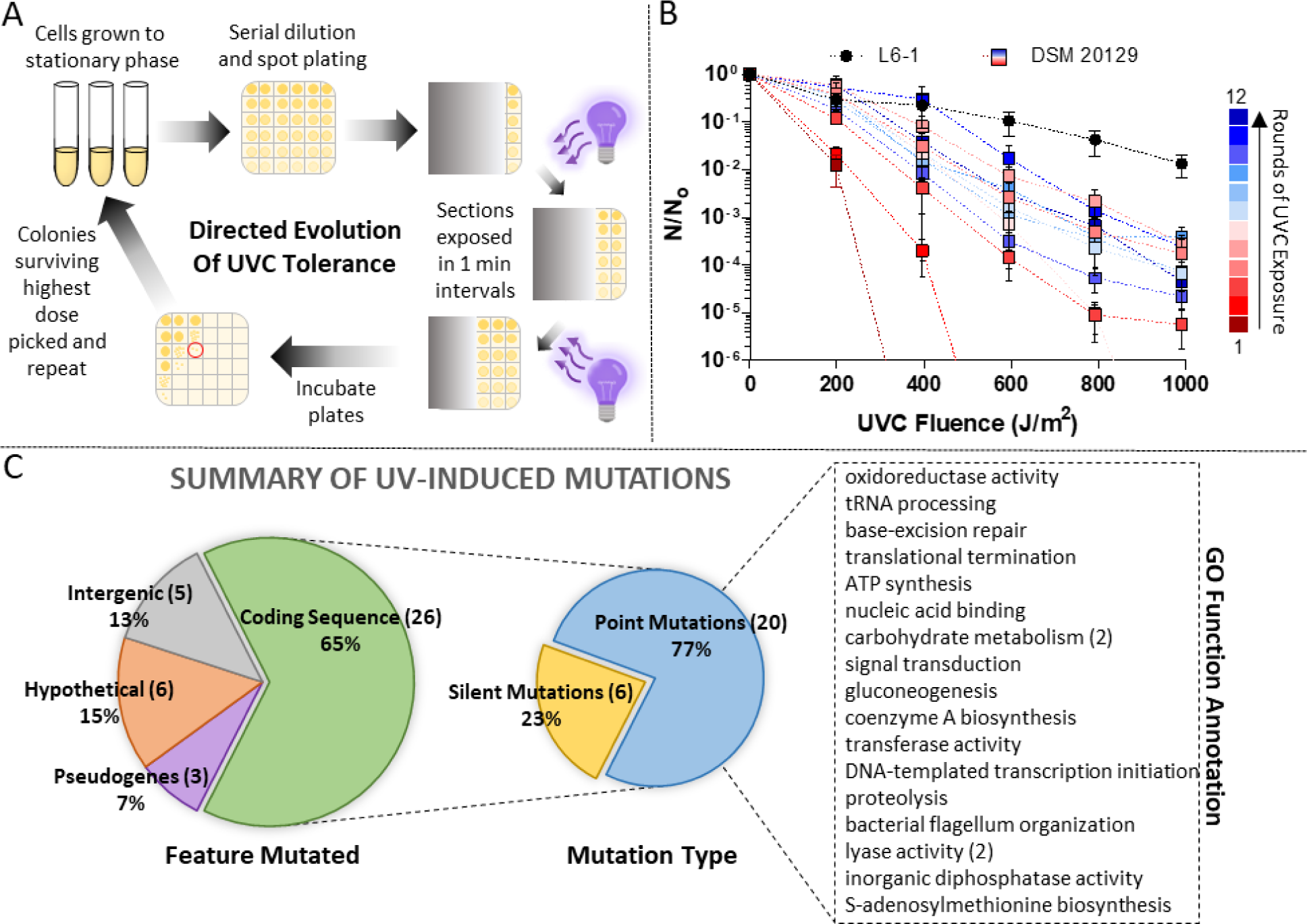
Directed evolution of UVCR tolerance in strain DSM 20129. **(A)** Schematic depiction of the directed evolution method used to improve UVCR tolerance in strain DSM 20129. The details of the procedure are descried in the text. **B)** UVCR tolerance of DSM 20129 after each round of UVCR exposure as compared to isolate L6-1, measured as the number of surviving CFUs (N) at each UVCR dose divided by total number of CFUs in the unexposed control (N_0_). Data shown are the average of three independent replicates. Error bars represent SEM. **C)** Summary of the mutations acquired during the directed evolution process. Mutation type (smaller pie chart) refers only to the mutations occurring in coding sequences (green slice in larger pie chart). Gene Ontology (GO) function annotations were determined for proteins that acquired amino acid point mutations in their coding sequence (blue slice in smaller pie chart).

Whole genome sequences were obtained for five evolved strains of DSM 20129 from the directed evolution experiment and where an increase in UVCR tolerance was observed (Table 2), and the mutations acquired were identified using the Breseq mutation analysis pipeline (40). In total, 40 mutations were identified in the most tolerant strain (DSM-9.3.3), with five occurring in intergenic regions, six in hypothetical genes, three in pseudogenes, and 26 in protein coding genes (Fig. 4C). Of the 26 mutations in protein coding genes, six are silent and 20 are nonsynonymous mutations (Table 2; Table A7) of proteins that participate in a wide range of reactions (Fig. 4C). The most notable are involved in DNA repair (uracil DNA glycosylase and DNA photolyase), cellular redox homeostasis (NAD(P)/FAD-dependent oxidoreductase in the thioredoxin reductase family), and the stress response (cold shock protein).

**Table 2:**
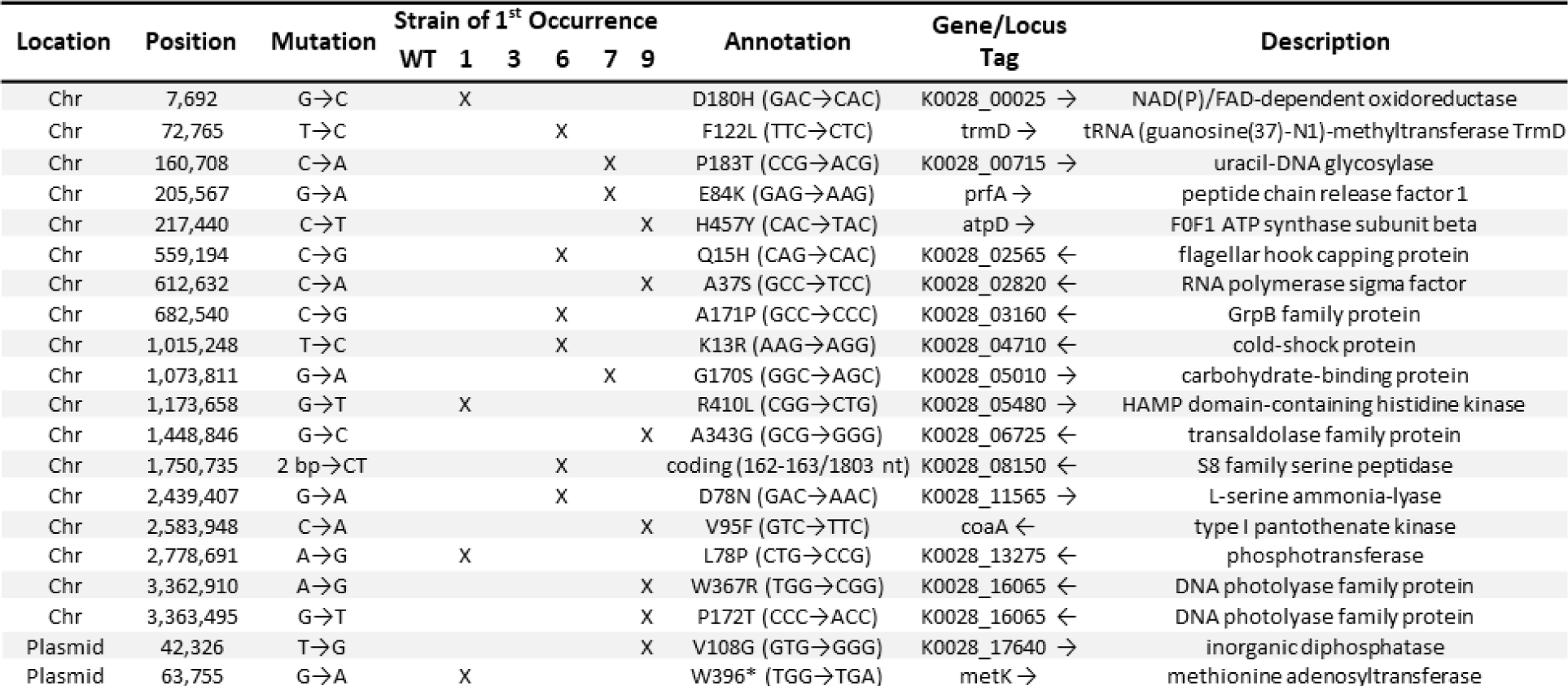
Mutations acquired in strain DSM 20129 during the directed evolution process.

### Intracellular ROS Concentrations After UVCR Exposure

Given the indirect effects of UVR exposure, we determined the concentration of ROS in the cells to assess if ROS detoxification may play a role in explaining the tolerances observed. ROS concentration before and after exposure to UVCR was monitored in isolate L6-1, the DSM 20129 parent strain, and the most UVCR-tolerant evolved strain (DSM-9.3.3) using the free radical sensing fluorescent probe H_2_DCFDA. ROS concentration did not significantly increase in L6-1, DSM 20129, or DSM-9.3.3 at any of the UVCR doses tested (up to 1,980 J m^-2^; Fig. 5A). Identical observations were made in experiments with *D. radiodurans* R1 (Fig. 5A), which is well known for its capacity to resist and detoxify oxidative stress-generating agents (41–43). In stark contrast, ROS concentrations in *E. coli* approximately doubled after each ∼1,000 J m^-2^ of UVCR exposure (Fig. 5A).

**Fig. 5.**
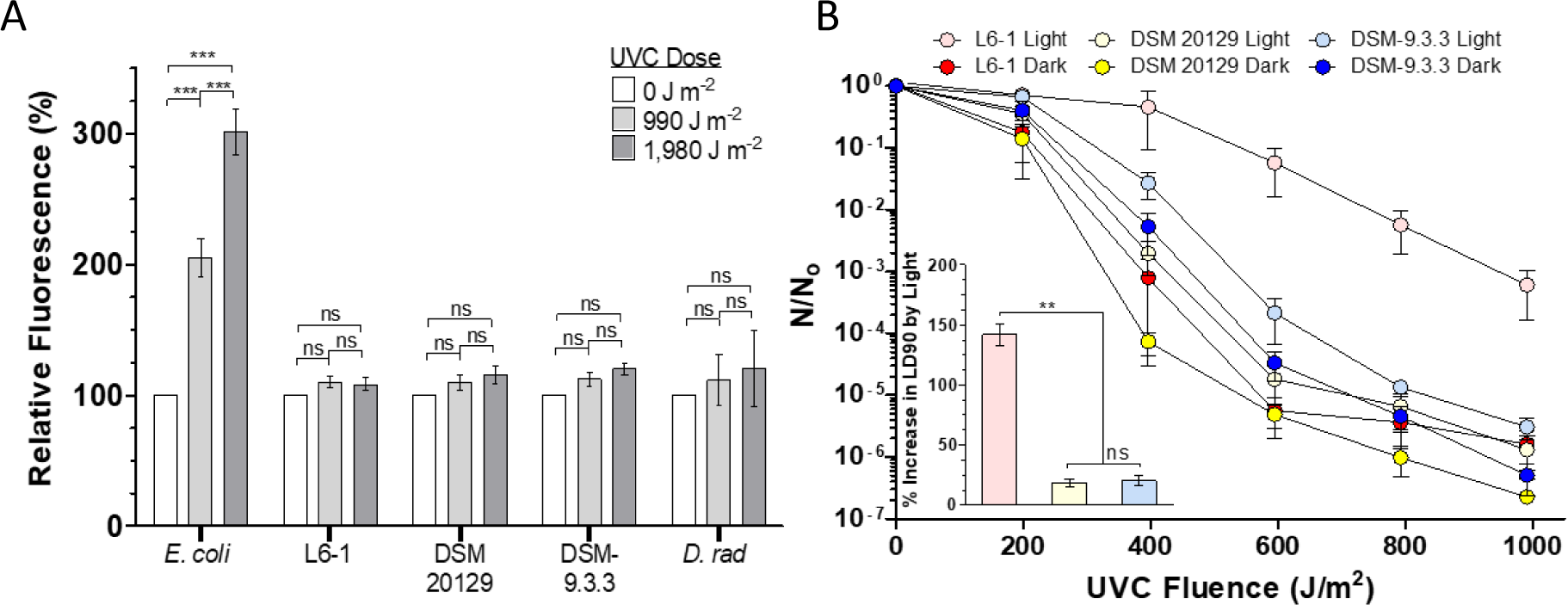
Cellular ROS concentrations and photolyase activity determination in UVCR treated cells. **A)** Intracellular ROS concentrations were quantified in the stratospheric isolate (L6-1), the DSM 20129 wild-type parent (DSM 20129), and UV-evolved strain (DSM-9.3.3) using the free radical probe H_2_DCFDA. *E. coli* MG1655 and *D. radiodurans* R1 were used as controls and for comparison. The average of three independent replicates with error bars representing SEM are plotted. **B)** UVCR survival of strains under dark repair and photoreactivating (light) conditions. Inset: Percent increase in LD_90_ after 1 h incubation in white light vs. dark after UVCR exposure. The data plotted are averages of four independent replicates with error bars representing SEM. (ns = not significant; ** = P-value < 0.01; *** = P-value < 0.001; one-way ANOVA and Tukey’s post-test).

### *In vivo* Assessment of Photolyase Activity

The contribution of photolyase to UVCR tolerance was investigated using an *in vivo* photorepair assay. Light activates photolyase for the repair of DNA photoproducts (44); therefore, its relative activity can be determined by examining the survival rate of UVCR exposed populations that recover under white light in comparison to those kept in the dark. Consistent with previous studies (45–47), post-UVCR exposure to white light increased the survival rate for all strains (Fig. 5B). Based on the interpolated LD_90_ values, there was not a significant difference in survival for DSM 20129 and DSM-9.3.3, indicating that the mutations incurred in the photolyase gene did not significantly affect DNA repair activity under the conditions tested (Fig. 5B inset). Photoreactivation of L6-1 increased the LD_90_ ∼140%, which is significantly higher (P-value = 0.004; one-way ANOVA) than the ∼19% increase observed for the DSM strains (Fig. 5B inset).

To determine if the second photolyase gene possessed by strain L6-1 contributes to its superior photorepair capacity, we aligned the sequences to known CPD and (6-4) photolyases. The CPD photolyases from *E. coli* (EcPhrB) and *Agrobacterium fabrum* (AfPhrA) align well with one of the photolyases found in L6-1 (*phr2*: KM842_RS02105) and the photolyase from DSM 20129 (*phr*: K0028_16065) (Fig. A1A). However, the additional photolyase in L6-1 (*phr1*: KM842_RS01465) is more closely related (∼37% identity) to the bacterial (6-4) photolyases of *A. fabrum* (AfPhrB), *Cereibacter sphaeroides* (CsCryB), and *Vibrio cholerae* (Vc(6–4) FeS-BCP), and the active site residues determined for AfPhrB are largely conserved in all four species (Fig. A1B). Furthermore, the protein structural prediction algorithm AlphaFold2 was used to model 3D structures for the L6-1 and DSM 20129 photolyases with a high degree of confidence (Fig. A2A-C). 3D structural alignments with the *E. coli* CPD photolyase (EcPhrB, PDB ID: 1DNP) show high similarity with the L6-1 Phr2 and DSM 20129 Phr, with a template-modeling score (TM-score) of 0.87 for both proteins (Fig. A3A). The L6-1 Phr1 shows high similarity with the *A. fabrum* (6-4) photolyase (AfPhrB, PDB ID: 5KCM) with a TM-score of 0.89, further supporting its proposed function as a bacterial (6-4) photolyase (Fig. A3B).

To assess the contributions of *phr1* and *phr2* to survival under UVCR exposure, we cloned them into expression vectors for heterologous expression in *E. coli* BL21 cells. Wild type BL21 cells were rapidly inactivated upon UVCR exposure, with an LD_90_ of 16 J m^-2^ under photoreactivating conditions (Fig. A4A). Overexpression of the native *E. coli* CPD photolyase (EcPhrB) significantly increased the LD_90_ by more than 230% during photoreactivation (Fig. A4B). Photoreactivation by the putative CPD photolyases from the *Curtobacterium* strains (L6Phr2 and DSMPhr for L6-1 *phr2* and DSM 20129 *phr*, respectively) also improved the survival of BL21, increasing the LD_90_ by 330% and 270%, respectively (Fig. A4B). In contrast, expression of the putative (6-4) photolyase from strain L6-1 (L6Phr1) alone did not significantly improve UVCR survival of BL21 cells (Fig. A4B). However, when the two L6-1 photolyases were co-expressed (L6Phr1 + L6Phr2), photoreactivation significantly improved survival over L6Phr2 alone (Fig. A4B) and increased the LD_90_ by an average of 400%.

## Discussion

There is a strong gradient of increasing biological stressors with altitude in the atmosphere (48, 49), and under the extreme environmental conditions in the stratosphere, prolonged exposure to UVR and desiccation are most important to limiting survival (18, 19, 50–61). The bacterial isolates studied (*Curtobacterium* sp. L6-1 and *Noviherbaspirillum* sp. L7-7A) were isolated from aerosols sampled at altitudes of 18 to 29 km ASL in New Mexico, USA and investigated because of their high tolerance to UVCR and desiccation (20). We tested close phylogenetic relatives of these isolates and found that they do not have extraordinary tolerance to UVCR (Fig. 1), providing an opportunity for a comparative analysis. UVR-induced cellular damage occurs directly through dimerization of pyrimidine bases in DNA, and indirectly, through the ROS generated by photosensitization mechanisms (31, 62). Accordingly, we hypothesized that higher tolerance to UVCR by isolates from the stratosphere is due to possessing DNA repair and antioxidant genes not found in their sensitive relatives.

### Comparative genomic evidence for UVR tolerance in Noviherbaspirillum sp. L7-7A

To our knowledge, UVR tolerance in the genus *Noviherbaspirillum* has not been previously reported or examined. Though we were unable to improve UVCR tolerance in the *Noviherbaspirillum* reference strains through directed evolution, comparative genomic analyses identified 12 genes in isolate L7-7A involved in DNA repair and antioxidant systems that were not present in the *Noviherbaspirillum* reference strain genomes (Fig. 3; Tables A5-A6): *nth2*, *nth3*, *uvrA2*, *recD*, *ligC*, *alkB*, *umuD*, *sbcC*, *sbcD*, *katE*, *mntH2*, and *pqiA2*. Previous studies have implicated a role for these genes in UVCR tolerance (Supplemental Discussion), and as such, any one (or combination) of these 12 genes could be responsible for isolate L7-7A’s high UVCR tolerance. However, genes with unknown functions (∼21% of ORFs in the L7-7A genome) cannot be ruled out as contributing to this phenotype.

### Genetic basis of UVR tolerance in the Curtobacterium strains

Previous studies have documented high UVR tolerance in members of the *Curtobacterium*, implying that it may be a common phenotype in this genus. Sundin and Jacobs (63) examined UVCR tolerance in bacteria isolated from the phyllosphere of field-grown peanut plants and found that *Curtobacterium* strains represented the largest proportion (∼43%, n=213) of those with UVCR minimum inhibitory doses (MID_c_) greater than 150 J m^-2^. Highly UVCR resistant strains of *Curtobacterium* have also been isolated from desert rock varnish, with 17% remaining viable after a UVCR dose of 220 J m^-2^ (64). Our results are consistent with these studies, with even the most UVCR “sensitive” *Curtobacterium* strain (DSM 20129) having an LD_90_ of ∼100 J m^-2^. Notably, *Curtobacterium* sp. L6-1 has a UVCR LD_90_ of 470 J m^-2^, which is more than double of that reported for other members of this genus and near survival rates reported for *D. radiodurans* (660 J m^-2^; (65)). In general, a species’ tolerance to UVR approximates natural exposure (24), which makes high tolerance to UVCR a surprising phenotype for populations to maintain in surface environments. However, this phenotype would be highly relevant for enabling survival and persistence at high altitudes in the atmosphere (20).

Comparative genomic analyses with DSM 20129 showed that strain L6-1 possesses seven additional genes related to DNA repair and antioxidant systems (Fig. 2; Tables A3-A4): *nei2*, *uvrD2*, *pcrA*, *phr1*, *katA*, *perR*, and *nrdH2*. Endonuclease VIII (*nei2*) is a bifunctional DNA N-glycosylase and abasic (AP) site lyase involved in the BER pathway. It initiates DNA repair by recognizing and removing oxidative base lesions and cleaving the phosphodiester backbone of the resulting AP site (66, 67). The UvrD DNA helicase II protein unwinds DNA in the 3′-5′ direction and plays a critical role in recombination, NER, and MMR (68–70). PcrA is a homolog of the UvrD helicase found in Gram-positive bacteria and has also been shown to participate in repairing UVR-induced DNA damage in *Bacillus* species (71).

Glutaredoxins (Grx) and glutathione (GSH), the main nonenzymatic antioxidants in Gram-negative bacteria, are generally absent in Gram-positive species. Many actinobacterial species are known to possess an alternative antioxidant system involving mycothiol (MSH), a low molecular weight thiol (72). Both L6-1 and DSM 20129 possess homologs for the *mshA-D* operon of the MSH biosynthesis pathway, and thus, MSH may serve as an important antioxidant in these strains. An additional copy of the gene encoding glutaredoxin-like protein NrdH in strain L6-1 is notable as NrdH has been shown to perform similar functions as the Grx/GSH system and is reducible by thioredoxin reductase (73, 74), an enzyme ubiquitous in all domains of life. In the GSH-lacking bacterium *Corynebacterium glutamicum*, NrdH enhances oxidative stress resistance (75), suggesting the second *nrdH* gene in L6-1 could play a similar role.

### Cross-tolerance to environmental stress

Desiccation- and radio-tolerance phenotypes frequently co-occur in extremophilic bacteria and archaea (76–84). Given that the acute doses to which radiotolerant species are resistant greatly exceed natural ionizing radiation sources on Earth, there is no evolutionary basis for ionizing radiation resistance to arise through natural selection. Studies have demonstrated a genetic linkage between desiccation- and ionizing radiation-resistance in *D. radiodurans* R1, supporting the hypothesis that DNA repair mechanisms evolved to compensate for water loss are also highly effective at repairing the similar pattern of DNA damage produced by exposure to ionizing radiation (85). Similarly, there is no obvious fitness benefit to having high UVCR resistance in any modern environment on Earth besides the stratosphere. However, since water stress is a very common phenomenon in the biosphere, the genetic and biochemical mechanisms that enhance survival to desiccation (e.g., ROS detoxification) may also provide tolerance to UVCR. Further, when L6-1 and L7-7A are desiccated (25% RH) and exposed to UVCR, their LD_90_s are 1200 J m^-2^ and 580 J m^-2^, respectively (20). These rates of survival are 2-to 3-fold higher than those observed when water has not been actively removed from the cells (Table A2) and highlights another protective effect that desiccation resistance can provide against indirect effects of UVR exposure.

In addition to possessing efficient pathways for repairing DNA damage, the ionizing and UVCR ‘resistome’ also includes mechanisms to physiologically detoxify ROS generated during exposure to these high-energy wavelengths. ROS produced when cells are exposed to UVR or desiccated can oxidize lipids and proteins (86, 87), as well as react with many other cellular constituents to inflict damage. ROS detoxification is known to have a role in bacterial desiccation resistance (87) and radiosensitive strains of *D. radiodurans* are more susceptible to oxidative damage than radiotolerant strains (88). Oxidative stress is a major contributor to the damage caused by desiccation and radiation; therefore, efficient antioxidant systems are likely to be important traits of xero-and radio-tolerant species. Our experiments indicate the presence of efficient ROS detoxification mechanisms in L6-1 and DSM 20129, with neither showing any significant increases in intracellular ROS concentrations after UVCR exposure (Fig. 5A). This implies other members of the *Curtobacterium* could also possess this trait, a contention supported by a study that exposed phyllosphere communities to the highly oxidative compound ozone (5,000-10,000 ppb) and found that the relative abundance of *Curtobacterium* taxa increased by ∼4-fold, implying a high tolerance of ozone (89). Consequently, it is tempting to speculate on the role that microbial antioxidant mechanisms may play during stratospheric transport beyond enhancing microbial tolerance to UVR and desiccation, as the concentration of ozone at altitudes of 15 to 30 km ASL (∼1,000 to 8,000 ppb) is ∼1000-fold higher than air near the surface (90).

### Photoprotection strategies in Curtobacterium

The comparative genomics analysis and directed evolution experiments implicated photolyase family proteins in providing increased UVCR tolerance in the *Curtobacterium* strains (Figs. 2 and 4C; Table 2). In bacteria, UVR-induced DNA dimers are repaired through direct reversal by photolyases or the lesion is removed via the NER pathway. NER requires the coordinated action of multiple proteins for incision, unwinding, and excision of a damage-containing segment of DNA, followed by resynthesis and ligation of the excised nucleotides (22, 29, 91). However, because this is an energy-intensive process, a more measured response may be necessary if a generous supply of energy is not available to the cell. For instance, direct reversal of DNA photoproducts by photolyases requires a single protein and light. Given the opportunity to function, the selective advantages of the photolyase repair system would make it a trait favorable to any species in an energy limited, high UVR flux environment, including the high atmosphere.

Photolyases catalyze the direct reversal of pyrimidine dimers in DNA when activated with near-UV/blue light in a process called photoreactivation (44). Two major classes of DNA photoproducts are produced by UVR: cyclobutane pyrimidine dimers (CPD) and pyrimidine (6-4) pyrimidone photoproducts (6-4PP). Direct repair of these lesions is catalyzed by photolyases specific to each lesion type (44, 92). Though CPD photolyases are widespread amongst bacteria, only three bacterial (6-4) photolyases have been characterized to date: PhrB of *Agrobacterium fabrum* (45, 93) (formerly *Agrobacterium tumefaciens* (94)), CryB of *Cereibacter sphaeroides* (95, 96) (formerly *Rhodobacter sphaeroides* (97)), and Vc(6–4) FeS-BCP of *Vibrio cholerae* (98). However, phylogenetic comparison with photolyase genes in the eukaryotic cryptochrome/photolyase family suggests that (6-4) photolyases are more prevalent in prokaryotes than initially thought (93). Sequence alignments and predicted structural models of the L6-1 photolyases (Fig. A1) suggest it possesses a putative CPD photolyase (*phr2*: KM842_RS16065) and (6-4) photolyase (*phr1*: KM842_RS01465). Strain L6-1 has higher photorepair capacity than strain DSM 20129 (Fig. 5B), which only possesses a CPD photolyase.

To demonstrate functionality of the putative 6-4PP and CPD photolyases of L6-1, *phr1* and *phr2*, respectively, were heterologously expressed in *E. coli* and their effects on its UVCR tolerance were examined. Although expression of *phr1* did not increase UVCR tolerance over that observed in the control, expression of *phr2* increased survival by ∼325%, while co-expression of *phr1* and *phr2* increased the survival rate even further (Fig. A4). These results are likely due to the larger number of CPD lesions (75-90% of total) upon UV exposure as compared to 6-4PP generated (22, 99–101). If further study shows the L6-1 *phr1* to be a bona fide (6-4) photolyase, its activity alone might be expected to have a negligible effect on cell survival since the overwhelming amount of DNA damage would be CPD dimers. However, when *phr1* and *phr2* are co-expressed, the repair of both CPD and 6-4PP dimers has a synergistic effect on survival.

### Conclusion

We investigated the genetic basis for extreme UVCR resistance phenotypes in bacteria recovered from stratospheric air masses (20), where high UVCR fluxes and low water availability represent endmember values for these variables in the biosphere. Recognizing the genetic underpinnings for the expressed characteristics mitigating damage from UVCR was facilitated by the rich history of experimental work that has characterized the biochemistry and molecular biology of DNA repair processes in model organisms. In fact, the genomes of the proteobacterial and actinobacterial strains studied encode genes for most, if not all, of the typical complement of DNA repair proteins documented in *E. coli* and many other bacteria. This indicates that strains L7-7A and L6-1 operate their canonical DNA repair pathways in a manner more effective than other species and/or that survival to UVCR is enhanced through alternative mechanisms. For microbial species surviving long distance dispersal as aerosols in the atmosphere, efficient DNA repair systems, such as photoreactivation, and mechanisms that evolved to cope with water stress are likely to be valuable for enduring the high UVR fluxes that intensify with altitude.

## Materials and Methods

### Bacterial Strains and Culture Conditions

The bacterial strains used in this study are listed in Table A1. *Curtobacterium* sp. L6-1 and *Noviherbaspirillum* sp. L7-7A were isolated from aerosols collected using a helium balloon payload that sampled at altitudes of 18 to 23 km and 24 to 29 km ASL, respectively, near Ft. Sumner, New Mexico in 2013 (32). Reference strains phylogenetically related to isolates L6-1 and L7-7A and that were identified as available in culture collections were obtained from the German Collection of Microorganisms and Cell Cultures (DSMZ) and the United States Department of Agriculture (USDA) Agricultural Research Service Culture Collection (NRRL). These included *Curtobacterium flaccumfaciens pv. flaccumfaciens* (DSM 20129), *Mycetocola reblochoni* (LMG 22367), *Plantibacter flavus* (DSM 14012T), and *Noviherbaspirillum soli* (SUEMI10). *Noviherbaspirillum denitrificans* (TSA40) and *Noviherbasprillum autotrophicum* (TSA66) were kindly provided by Dr. Satoshi Ishii of the University of Minnesota. Unless otherwise noted, the actinobacterial and betaproteobacterial strains were cultured aerobically at 30°C with vigorous shaking in LB (10 g L^-1^ tryptone, 5 g L^-1^ yeast extract, 10 g L^-1^ NaCl) and R2A (Difco cat. no.: 218262) media, respectively. When required, media were amended with antibiotics at the following concentrations: spectinomycin (50 µg mL^-1^) & carbenicillin (100 µg mL^-1^).

### UVCR Survival Assays

Cultures were grown in 5 mL of liquid media to stationary phase, and six serial dilutions of this material were prepared in 10 mM MgSO_4_. Ten μL of each dilution was transferred to six sections of a square plate containing agar-solidified growth media. Portions of the plate were then covered with aluminum foil such that sections uncovered sequentially during 1 min interval exposures were provided with controlled doses of UVCR (GE G36T5 UVC Light Bulb; λ = 254 nm; 3.3 W m^-2^) that were 198, 396, 594, 792, and 990 J m^-2^. The UVCR dose was measured using a digital radiometer (Solar Light Company, Inc., Glenside, PA). The longest exposure for the populations was 5 min. and the section of the plate that remained covered during the experiment (i.e., was not exposed to UVCR) served as the control (N_0_). The cultures were incubated at 30°C for 24 to 48 h and the number of colony forming units (CFU) surviving each UVCR dose (N) was used to calculate the surviving fraction, expressed as N/N_0_. Survival rates are reported as the mean and SEM for three biological replicates.

### Whole Genome Sequencing and Comparative Genomics

Whole genome sequences for *Noviherbaspirillum sp.* L7-7A, *Curtobacterium sp.* L6-1, and *Curtobacterium flaccumfaciens* strain DSM 20129 were obtained as described previously (33). Briefly, stationary phase cultures were pelleted, frozen (-70°C), and shipped to SNPsaurus (Eugene, OR) for DNA extraction, library preparation, sequencing, and genome assembly using the PacBio Sequel II sequencing platform followed by de novo genome assembly using the Flye v2.7 assembler (102). The assembled genomes were submitted to GenBank (L7-7A: <u>JAHQRJ01</u>; L6-1: <u>CP076544.1</u>; DSM 20129: <u>CP080395.1</u>/<u>CP080396.1</u>) and annotated using the NCBI Prokaryotic Genome Annotation Pipeline (PGAP) (103).

Average nucleotide identity (ANI) and average amino acid identity (AAI) among the isolates and reference strains were calculated using the ANI and AAI calculators from the Kostas lab (http://enve-omics.ce.gatech.edu/) (104, 105). Functional comparisons were made with the Rapid Annotation using Subsystem Technology (RAST) server and the SEED Viewer (106–108). Further comparisons were made by searching for homologs of known DNA repair and ROS detoxification proteins from model organisms in the L6-1, DSM 20129, L7-7A, TSA40, and TSA66 genomes using the blastp algorithm from NCBI’s Basic Local Alignment Search Tool (BLAST) (109). Query protein sequences were obtained from the UniProt database for *Escherichia coli* strain K12, *Bacillus subtilis* strain 168, and *Mycobacterium tuberculosis* strain ATCC 25618 / H37Rv (110). Reciprocal best hits were considered homologs if the bit score was >50, E-value was <1e-5, and query coverage was >50% (111). Differences in gene content among the isolates and the reference strain relatives were visualized using the BLAST Ring Image Generator (BRIG) (112).

### Directed Evolution and Mutation Analysis

The survival of strain DSM 20129 to UVCR exposure was determined as described above. Triplicate colonies surviving the highest UVCR dose were selected, grown separately in liquid media, exposed to UVCR (as described in the section “UVCR Survival Assays” above), and the process was repeated a total of 12 times (Fig. 4A). The populations from each cycle of exposure were archived by freezing aliquots of the cultures in 25% glycerol and storing at -70°C. Genome sequences were obtained for select strains throughout the directed evolution process using the Illumina NextSeq 2000 sequencing platform (Microbial Genome Sequencing Center, Pittsburgh, PA) with 2×150 bp reads. Sequencing reads were quality filtered and adaptors trimmed using the Trim Galore script (113), followed by mapping to the reference genome and mutation identification with the Breseq mutation analysis pipeline (40). The ancestral parent strain was sequenced and used as a control to correct the reference genome before comparison with the evolved strains. Gene Ontology (GO) functional annotations for proteins incurring mutations throughout the directed evolution process were obtained using Blast2GO (114).

### ROS Quantification Assays

The free radical probe 2’,7’-dichlorodihydrofluoresceindiacetate (H_2_DCFDA) (Biotium, Fremont, CA) was used to quantify ROS. Stationary phase cultures of *E. coli* MG1655, *D. radiodurans* R1, isolate L6-1, the wild-type parent strain DSM 20129, and UV-evolved strain DSM-9.3.3 were diluted to OD_600_ 1.0 in 10 mL of media. Cells were washed with 10 mM potassium phosphate buffer (pH 7.0) and incubated for 30 min at 25°C in the same buffer containing 10 μM H_2_DCFDA. Subsequently, cells were washed and resuspended in 10 mL potassium phosphate buffer (10 mM; pH 7.0), transferred to 60×15 mm petri plates, and exposed to UVCR doses of 0, 990, or 1,980 J m^-2^ while stirring the cell suspensions at 400 rpm with a magnetic stir bar. The exposed cell populations were then washed, resuspended in fresh buffer, and disrupted by vortexing with lysing matrix B (MP Biomedicals). Cellular debris was removed by centrifugation for 10 min at 5,000×g, and fluorescence intensity in the supernatant was measured using a multi-well plate reader (Molecular Devices SpectraMax M3; Exc. 490nm, Emm. 519nm). The amount of fluorescence observed was normalized per mg of protein, as determined by the Pierce BCA Protein Assay Kit (Thermo Scientific) and compared relative to data from the unexposed control (i.e., time zero is 100% fluorescence intensity). Values reported are the average of three biological replicates and the error bars indicate SEM.

### Photolyase Activity Assays and Sequence/Structural Comparisons

To assess *in vivo* photorepair, isolate L6-1, the DSM 20129 founder strain, and UV-evolved strain DSM-9.3.3 were grown to stationary phase and exposed to UVCR using the same method as described for the UVCR survival assays. After exposure, half of the cultures were immediately placed in the dark and the other half were photoreactivated for 1 h under a daylight lamp (GE F20T12-C50-ECO 20W Chroma 50 5000K Ecolux Daylight Light Bulb). All of the cultures were subsequently incubated in the dark at 30°C until colonies formed. Survival rates under dark and light conditions were calculated, and the lethal dose reducing the population by 90% (LD_90_) was determined by fitting the survival curves to an exponential decay model. The percent increase in LD_90_ by photoreactivation was calculated using the equation: 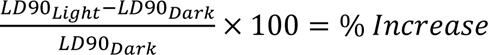. Results shown represent the average of four biological replicates with error bars representing SEM.

## Acknowledgements

This work was supported by funding from the University of Florida Institute of Food and Agricultural Sciences and awards from the National Aeronautics and Space Administration (NASA) Exobiology Program (80NSSC21K0486) and the University of Central Florida NASA Florida Space Grant Consortium and Space Florida (NNX15034). B.C.C. is indebted to John R. Battista for enduring countless discussions on bacterial radio- and desiccation-tolerance. Satoshi Ishii of the University of Minnesota kindly provided strains for the analysis. We thank Madison Drum for assistance with the directed evolution experiments. We also thank T. Gregory Guzik, Noelle Bryan, Doug Granger, Michael Stewart, and the many other dedicated members of the 2009-2013 ACES and HASP balloon programs at Louisiana State University for their tireless efforts in developing/testing the aerosol sampling payloads, conducting the missions (and their recovery, no matter where it landed!), and performing the initial research that made this study possible.

